# Endogenous oxytocin reconfigures brainwide network dynamics toward salience-related states

**DOI:** 10.64898/2026.09.22.753403

**Authors:** A. Elizabeth de Guzman, Caterina Montani, Andrew Hayward, Sara Migliarini, Giovanni Morelli, Daniel Gutierrez-Barragan, Alberto Galbusera, Filomena Alvino, Ludovico Coletta, Yongsoo Kim, Stefano Panzeri, Laura Cancedda, Massimo Pasqualetti, Alessandro Gozzi

## Abstract

Oxytocin (OXT) can exert diverse effects across social, affective and motivational domains, yet its impact on brain function at the systems level remains unclear. Here, we combine chemogenetics with fMRI and electrophysiology in mice to investigate how endogenous OXT reshapes intrinsic large-scale brain network dynamics. Using mice constitutively expressing hM3Dq DREADD receptors in OXT-releasing hypothalamic neurons, we show that chemogenetic stimulation of OXT neurons elicits widespread increases in cerebral blood volume, with prominent effects in fronto-striatal regions and dorsal hippocampal areas. This broad functional response is accompanied by a network-specific reconfiguration of brain-wide functional connectivity, characterized by increased coupling between key components of the rodent social brain, and negative functional coupling between hippocampal and fronto-cortical regions. *In vivo* electrophysiology under matched conditions shows concordant changes in slow (<1 Hz) and δ-band coherence across frontal-thalamic and fronto-hippocampal circuits, revealing a plausible neural correlate of the observed connectivity effects. Notably, these changes are accompanied by a marked reconfiguration of the temporal architecture of fMRI network activity, with increased occupancy of co-activation states engaging salience-related regions and a concomitant suppression of sensory-dominated states. Together, these findings identify dynamic network-state reconfiguration as a systems-level mechanism through which endogenous OXT can tune functional communication across social-affective circuits.

## Introduction

Oxytocin (OXT) modulates a broad range of social, affective and motivational behaviors, including affiliation, parental responses, stress regulation and social vigilance (Yao & Kendrick, 2025). Across these domains, OXT effects are often subtle, multifaceted and strongly context dependent (J. A. Bartz et al., 2011). For example, depending on environmental cues and internal states, OXT can promote either prosocial or antisocial responses (J. Bartz et al., 2011; Declerck et al., 2010), and increase or suppress stress-related behaviors (Eckstein et al., 2014; Guzman et al., 2013). Several theoretical frameworks have been proposed to explain the highly heterogeneous and seemingly opposing behavioral effects of OXT, including the social salience hypothesis (Shamay-Tsoory & Abu-Akel, 2016) the approach/withdrawal hypothesis (Harari-Dahan & Bernstein, 2014), the neural circuit framework (Jurek & Neumann, 2018), and the allostatic theory (Quintana & Guastella, 2020). Importantly, these perspectives converge to suggest that the breadth and context dependence of OXT action may not be fully captured by projection-specific circuit mechanisms alone, but may also involve inter-areal coordination across distributed brain systems. However, whether endogenous OXT can engage such large-scale functional organization remains unknown.

Such a distributed mode of action is plausible, and consistent with the anatomical organization of the oxytonergic system. OXT is synthesized mainly in the hypothalamus, from which it is released either into the bloodstream or centrally through axonal projections to act on oxytocin receptors (OXTRs) (Stoop, 2012). Interestingly, the distribution of OXT projections only partially overlaps with the broader anatomical expression of OXTRs (Son et al., 2022), suggesting that endogenous OXT may not merely influence brain activity through directly innervated targets, but could also recruit distributed brain systems on a slower timescale through volume transmission (Parmaksiz & Kim, 2025). Human and animal imaging studies are consistent with this view, as exogenous (i.e., intranasal)) OXT administration has been shown to elicit widespread modulation of brain activity and functional connectivity in cortical and subcortical regions extending beyond the areas most strongly innervated by OXT (Galbusera et al., 2017; Martins et al., 2020; Pagani et al., 2020; Paloyelis et al., 2016). However, our understanding of the systems-level effects of OXT derives primarily from pharmacological studies in which the neuropeptide is administered exogenously. As a result, the brainwide substrates and large-scale functional effects of endogenously-released OXT remain largely unknown.

Here, we combine DREADD-based chemogenetics (Roth, 2016), fMRI and *in vivo* electrophysiology in the mouse to map the network-level substrates engaged by chemogenetically evoked OXT-neuron activation. We refer to this manipulation as “endogenous OXT release” to distinguish OXT signaling produced by chemogenetic activation of OXT-producing neurons, from that produced by exogenously administered peptide, as used in previous intranasal or systemic OXT studies. We find that endogenous OXT release robustly reconfigures connectivity across hypothalamic, fronto–thalamic and fronto– hippocampal circuits, alters interareal low-frequency synchrony, and biases spontaneous activity toward salience-related fMRI states. These results outline a network mechanism through which endogenous OXT can tune large-scale socio-affective circuits.

## Results

### Chemogenetic activation of OXT neurons evokes endogenous OXT release

To enable controlled endogenous OXT release, we leveraged a conditional hM3Dq knock-in mouse line carrying a double-floxed hM3Dq DREADD allele (Giorgi et al., 2017), which we crossed with Oxt-IRES-Cre animals (Wu et al., 2012), thereby generating mice constitutively expressing hM3Dq in OXT neurons (hereafter, OXT-hM3Dq). To probe the specificity of DREADD receptor expression, in a separate validation cohort we crossed Oxt-IRES-Cre mice with the Rosa26-tdTomato reporter line (Figure 1a). Immunostaining revealed extensive overlap between reporter expression and OXT immunoreactivity in neurons of the paraventricular nucleus (PVN) (Figure 1a), consistent with selective Cre-dependent recombination in OXT cells. We further characterized hM3Dq expression in OXT-hM3Dq mice using *in situ* hybridization for Oxt mRNA and immunohistochemistry for hM3Dq/mCherry in adjacent PVN sections **(Figure 1b)**. Cell counts revealed comparable numbers of OXT-positive and hM3Dq-positive cells across slices, thus corroborating the specificity of our manipulation (mixed-effects TOST for equivalence, ΔL = -10, p=0.03, ΔU = 10, p=0.04; **Supplementary Figure 1**).

**Figure 1.**
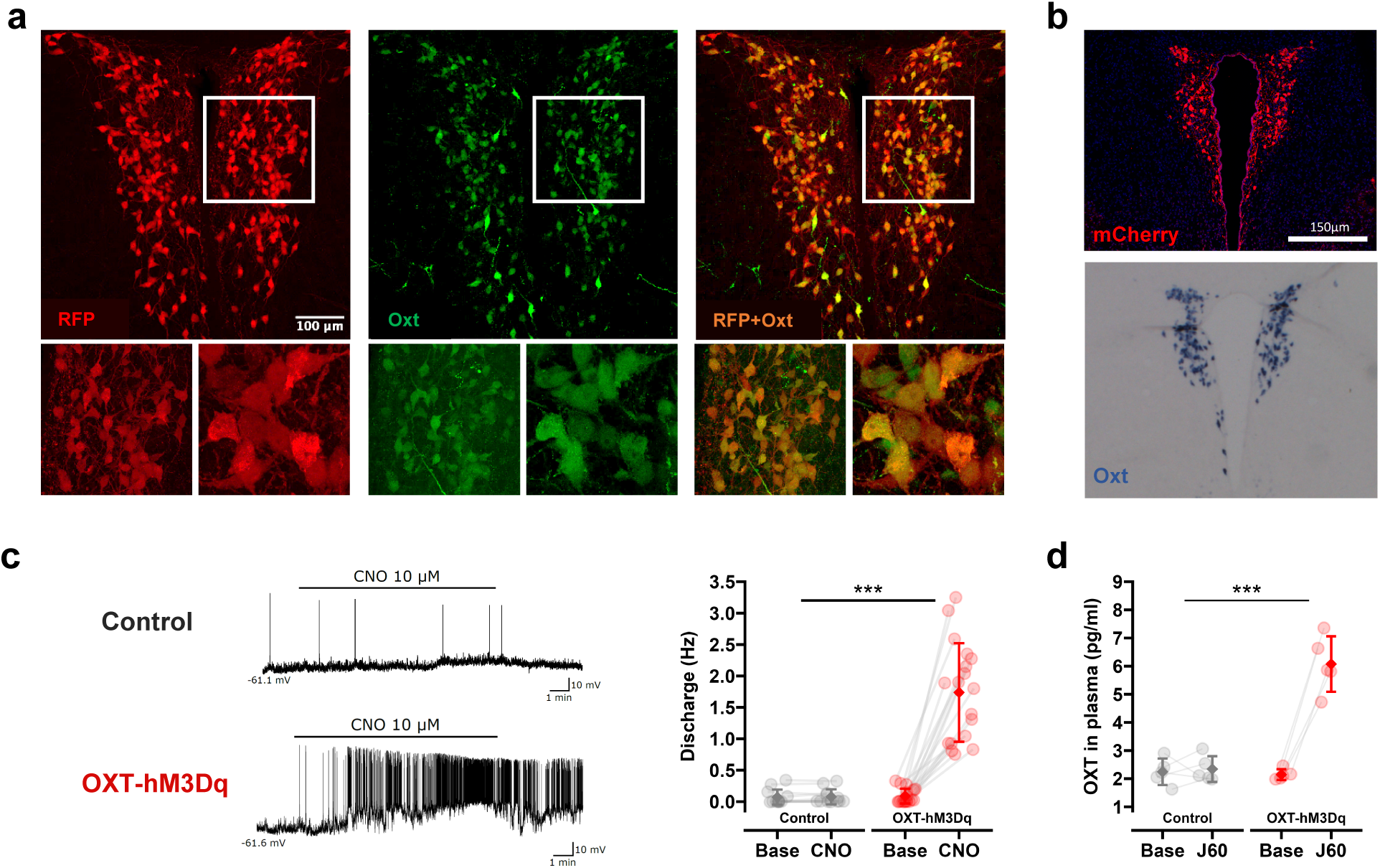
Validation of a chemogenetic model for endogenous OXT release. (a) Representative immunohistochemistry showing RFP expression (red) and OXT-immunoreactive neurons (green) in the PVN of OXT-Ires-Cre × ROSA26-tdTomato reporter mice. (b) Representative hM3Dq-mCherry immunoreactivity and Oxt mRNA expression detected by in situ hybridization in the PVN of OXT-hM3Dq mice. (c) Whole-cell current-clamp traces and quantification of firing frequency before and after CNO bath application in PVN neurons from OXT-hM3Dq (responding cells only, increase in frequency discharge >0.5 Hz) and control mice. Data are shown as mean ± 95% CI; treatment × genotype effect, linear mixed model, ***p < 0.001. See Supplementary Figure 3 for the analysis of all recorded PVN cells. (d) Plasma OXT levels measured before and after J60 administration in OXT-hM3Dq and control mice. Data are shown as mean ± 95% CI; treatment × genotype effect, linear mixed model, ***p < 0.001. PVN, paraventricular nucleus.

We next examined whether chemogenetic activation of hM3Dq receptors increased firing activity of PVN neurons in OXT-hM3Dq mice. Whole-cell recordings showed that bath application of the DREADD agonist clozapine-N-oxide (CNO, 10 μM) significantly increased firing in approximately 50 % (18/37) of the patched neurons recorded from OXT-hM3Dq mice, whereas wild-type PVN neurons showed no apparent CNO-evoked increase in firing (Figure 1c, linear mixed model treatment*genotype effect, Satterthwaite’s p<0.001). Importantly, these effects were observed in neurons with both parvocellular- and magnocellular-like electrophysiological profiles (Supplementary Figure 2). The effect of CNO remained significant also when all recorded PVN neurons were included, irrespective of their electrophysiological profile, or responsiveness to CNO (treatment × genotype interaction, linear mixed model, permutation p < 0.05; Supplementary Figure 3).

To determine whether *in vivo* chemogenetic activation of DREADD-expressing neurons resulted in endogenous OXT release, we measured plasma OXT levels in OXT-hM3Dq and control mice upon administration of DREADD agonists. Previous work showed that clozapine, a metabolite of CNO, can stimulate peripheral OXT release in wild-type rats (Uvnas-Moberg et al., 1992). In our *in vivo* chemogenetic experiments, we thus replaced CNO with the selective DREADD agonist JHU37160 (J60, (Bonaventura et al., 2019)). Accordingly, J60 administration did not increase plasma OXT levels in control mice, but elicited a robust rise in circulating OXT in OXT-hM3Dq animals (Figure 1d; ∼3-fold increase; linear mixed model treatment × genotype interaction, p < 0.001). This peripheral response falls within the lower physiological range reported in rodents (Grippo et al., 2007), and remains well below the larger neuroendocrine surges associated with parturition, suckling, or milk ejection (Grosvenor et al., 1986). Taken together, these results show that J60 administration in OXT-hM3Dq mice activates OXT-producing neurons and elicits endogenous OXT release.

### Endogenous OXT elicits widespread increases in cerebral blood volume

To map the brain-wide effects of endogenous OXT release, we used contrast-enhanced fMRI to quantify relative changes in cerebral blood volume (rCBV) produced by chemogenetic activation of OXT-producing neurons. Acute administration of J60 elicited widespread rCBV increases across multiple brain regions in OXT-hM3Dq mice (two-level mixed-effects analysis of pre vs post 30min COPE, |Z| > 2.1, cluster corrected p<0.05; Figure 2a). The temporal profile of this response revealed a gradual and sustained increase in rCBV signal, reaching maximal levels around 20-40 min after J60 administration (Figure 2b). Clusters of significant rCBV increases were observed in the hypothalamus, dorsal striatum, anterior cingulate and retrosplenial cortices (FDR corrected q < 0.1, Figure 2a, c, Supplementary Table 1). However, inspection of unthresholded functional maps (Figure 2a) and region-wise quantification across anatomically defined areas (Figure 2c) revealed that OXT-induced rCBV response was widespread, extending beyond the regions surviving statistical thresholding.

**Figure 2.**
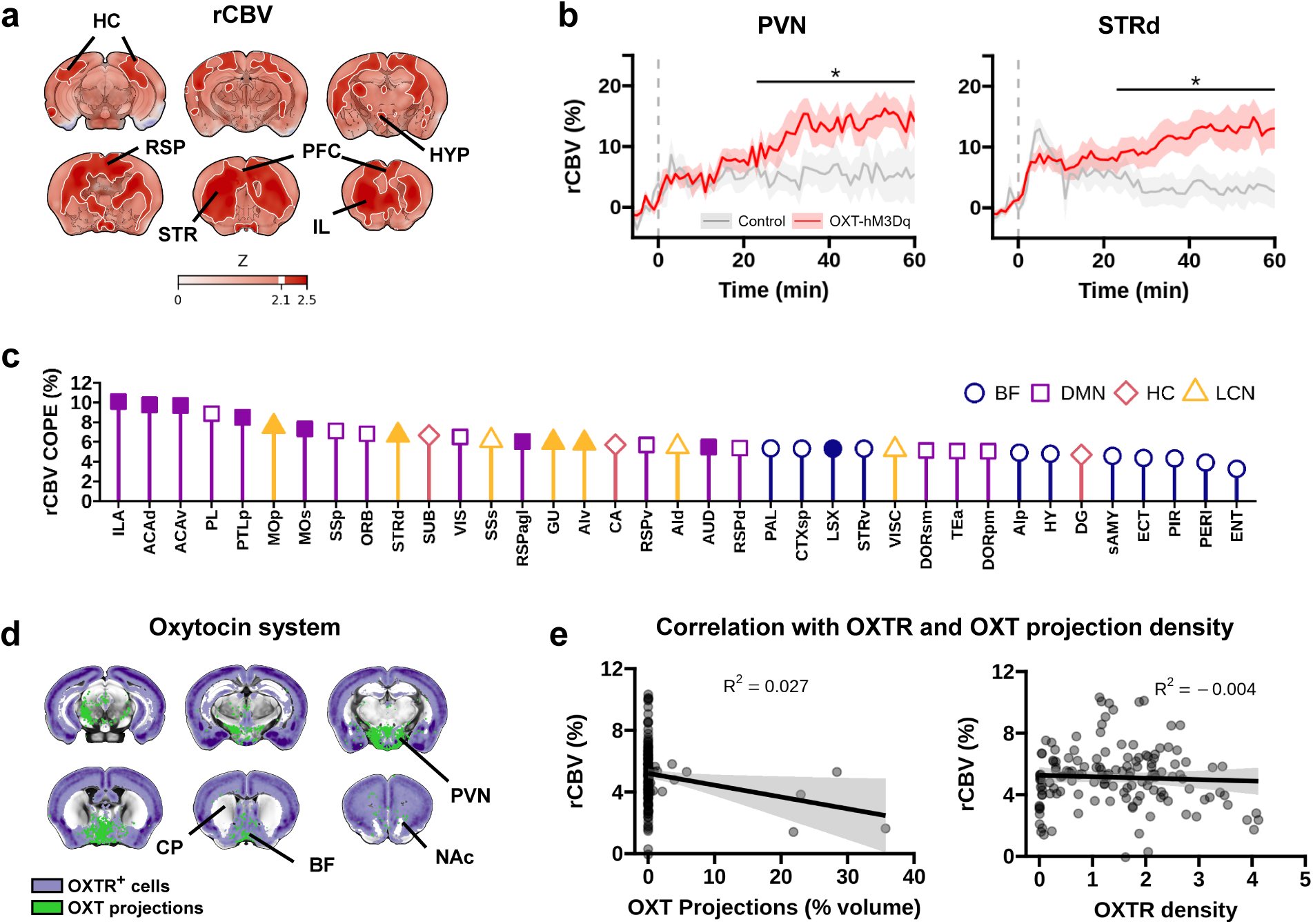
Endogenous OXT induces widespread increases in cerebral blood volume. (a) Statistical parametric maps (Z) showing relative cerebral blood volume (rCBV) increase produced by J60 administration in OXT-hM3Dq mice compared to control animals (|Z| > 2.1, cluster-corrected p < 0.05). (b) Temporal profile of rCBV (group mean ± 95% CI) following J60 injection (dotted line). *q < 0.1 for between-group differences in the estimated post-injection rCBV response, FDR-corrected across all examined regions. (c) Regional between-group differences in the estimated post-injection rCBV response (filled symbols: FDR-corrected q < 0.1 across regions, COPE: contrast of parameter estimates). (d) Oxytocin receptor (OXTR) density and OXT cell projection maps. Replotted from (Son et al., 2022). (e) Relationship between regional rCBV responses and OXT projection density or OXTR density across regions shown in (c). rCBV responses were not significantly predicted by OXT projection density (F_(1,70)_ = 2.6, p = 0.1) or OXTR density (F_(1,70)_ = 0.1, p = 0.7; linear regression). PVN, paraventricular nucleus; dSTR, dorsal striatum; BF, basal forebrain; HC, hippocampal network; DMN, default mode network; LCN, lateral cortical network.

Because our chemogenetic activation also increased peripheral OXT levels, we next asked whether the observed rCBV effects could be confounded by systemic cardiovascular responses (i.e., related to breakdown of cerebral autoregulation (Gozzi et al., 2007), rather than central functional activation). Under the same experimental conditions used for fMRI, however, chemogenetically evoked OXT release did not produce appreciable alterations in arterial blood pressure (Supplementary Figure 4). This finding argues against a confounding contribution of peripheral cardiovascular components to the mapped rCBV effects.

We finally asked whether the spatial distribution of the observed rCBV response could be explained by the known anatomy of the oxytocin system, as indexed by OXTR expression and oxytocinergic projection density (Son et al., 2022). Visual comparisons suggested only limited correspondence between the spatial pattern of OXT-induced rCBV effects, and these anatomical maps (Figure 2d). A representative example was the dorsal striatum, which showed a robust rCBV increase, despite very sparse OXTR expression and projection density. Consistent with these qualitative observations, the magnitude of the regional rCBV response did not significantly correlate with either OXTR expression map, or oxytocinergic projection density (F_(1,70)_ = 0.1, p = 0.7, F_(1,70)_ = 2.6, p = 0.1; Figure 2e). Together, these findings indicate that the brain-wide functional response to endogenous OXT release is not readily predicted by the anatomical distribution of OXTR expression or oxytocinergic projections.

### Endogenous OXT reconfigures brain-wide functional connectivity

The widespread rCBV response produced by endogenous OXT release raises an apparent paradox: how can such broad regional recruitment be reconciled with the selective circuit-level actions previously reported for OXT (Ferretti et al., 2019; Knobloch et al., 2012; Tan et al., 2019; Thirtamara Rajamani et al., 2024; Tsai et al., 2022)? One possibility is that, rather than acting solely through regional activation, OXT may also act by selectively reconfiguring functional communication across brain areas. To test this hypothesis, we mapped resting-state fMRI connectivity in OXT-hM3Dq and control mice before and after J60 administration.

First, we computed global fMRI connectivity, a voxel-wise measure of how strongly each voxel is functionally coupled to the rest of the brain (Pagani et al., 2025). This analysis revealed focal increases in global connectivity centered on the hypothalamus and basal forebrain in OXT-hM3Dq mice (t-test, |Z| > 2.1, cluster-corrected p < 0.05; Figure 3a), a finding consistent with the anatomical organization of the OXT system. Because global connectivity identifies where connectivity changes occur, but not which regions are specifically coupled to one another, we next mapped interregional fMRI connectivity across the whole brain using a functional parcellation (Coletta et al., 2020) (Figure 3b). Inspection of the resulting connectivity matrix revealed a non-uniform pattern of connectivity changes across multiple brain systems, involving both intra- and inter-network effects, rather than a uniform shift in connectivity strength.

**Figure 3.**
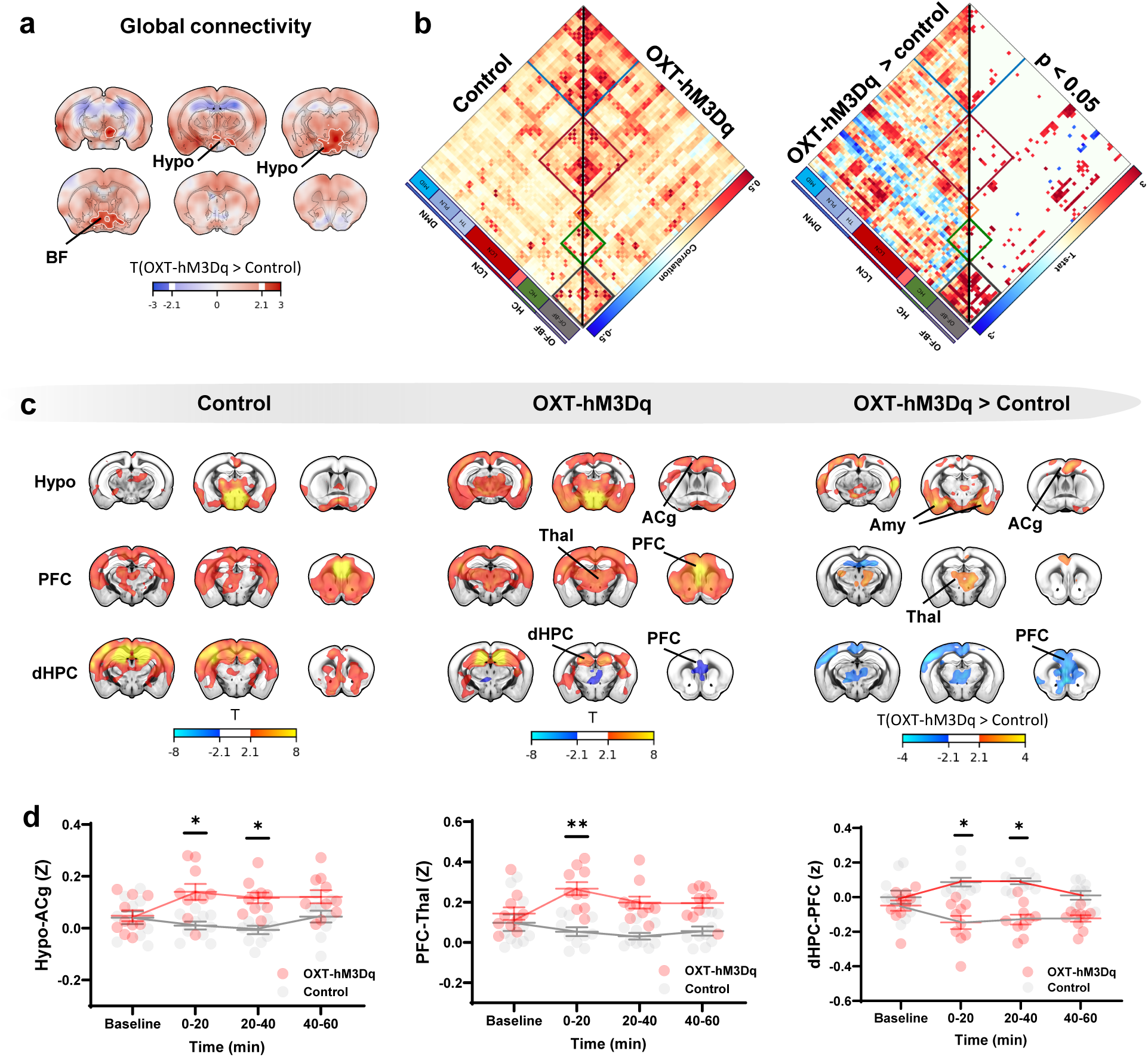
Endogenous oxytocin reorganizes large-scale functional connectivity. (a) Voxel-wise analysis of global functional connectivity (FC) showing increased hypothalamic and BF connectivity following J60 administration in OXT-hM3Dq mice relative to controls (|T| > 2.1, cluster-corrected p < 0.05, unpaired t test). (b) Left, group-averaged functional connectivity matrices for control (left half) and OXT-hM3Dq (right half) mice, with regions ordered by connectivity modules (Coletta et al., 2020). Right, between-group connectivity differences expressed as t-statistics, shown without thresholding (left half) and at p < 0.05, uncorrected (right half). (c) Seed-based correlation maps in control and OXT-hM3Dq mice, and corresponding between group connectivity difference map for hypothalamic, prefrontal, and dorsal hippocampal seed regions. Seed maps and group-difference maps are thresholded at |T| > 2.1, cluster-corrected p < 0.05. (d) Functional connectivity between selected region pairs across baseline and post-J60 time windows. Data are shown as group mean ± 95% CI; group × time interaction, linear mixed-effects model, *p < 0.05, **p < 0.01. Hypo, hypothalamus; BF, basal forebrain; PFC, prefrontal cortex; Thal, thalamus; dHPC, dorsal hippocampus; ACg, anterior cingulate cortex; Amy, amygdala.

We then used seed-based analyses to probe specific network components of this functional reconfiguration, focusing on systems of particular relevance to OXT signaling and its socio-affective effects (Froemke & Young, 2021), such as the hypothalamus, medial prefrontal cortex (PFC), and dorsal hippocampus, corresponding to the olfactory/basal forebrain (OF-BF), default mode (DMN), and hippocampal (HC) networks, respectively (Gutierrez-Barragan et al., 2022). This analysis revealed that endogenous OXT release elicited robust network-specific changes in functional connectivity. Specifically, following J60 administration, OXT-hM3Dq mice showed increased fMRI connectivity between the hypothalamus and both the amygdala and PFC, as well as between the PFC and thalamus, together with reduced fMRI connectivity between the dorsal hippocampus (dHPC) and fronto-thalamic areas (Figure 3c, t-test, |Z| > 2.1, cluster-corrected p < 0.05). Interestingly, the observed reduction in hippocampal-prefrontal connectivity was driven by the emergence of negative fMRI coupling between these regions in OXT-hM3Dq mice, but not in control animals (Figure 3d). Together, these results show that endogenous OXT release reconfigures functional connectivity across distributed brain systems and suggest that such selective interareal reorganization may provide a systems-level substrate for the circuit specificity of OXT action.

### Endogenous OXT reorganizes local electrophysiological activity and interareal synchrony

The observation of large-scale fMRI connectivity changes after endogenous OXT release raised the question of whether these signatures were accompanied by measurable electrophysiological changes within the implicated circuits. To address this question, we carried out *in vivo* electrophysiological recordings in the PFC, hippocampus, and mediodorsal thalamus (Supplementary Figure 5), three regions showing prominent functional connectivity changes in our fMRI analyses.

J60 administration in OXT-hM3Dq mice significantly increased local field potential (LFP) power across all three recorded regions (Figure 4a-b). This effect involved broad LFP power modulation across the probed frequency spectrum. In the PFC, the increase in LFP power was most prominent at slow and δ frequencies, whereas in the HPC it was more evenly distributed across frequency bands. In the thalamus, significant LFP power modulation was mainly detected at higher frequencies (two-sided Wilcoxon rank-sum tests, cluster-corrected p < 0.01; Figure 4b). Together, these findings show that endogenous OXT reorganizes local oscillatory activity across distributed brain regions. Multi-unit activity analyses revealed more heterogeneous effects. J60 did not significantly alter MUA rate in the PFC, whereas in the HPC it produced a marked separation between groups, with increased activity in OXT-hM3Dq mice and reduced activity in controls (genotype × time interaction, p = 1 × 10⁻⁶; Supplementary Figure 6). In the thalamus, a transient reduction in MUA rate in OXT-hM3Dq mice was observed, but this effect did not reach statistical significance (p = 0.135; Supplementary Figure 6).

**Figure 4.**
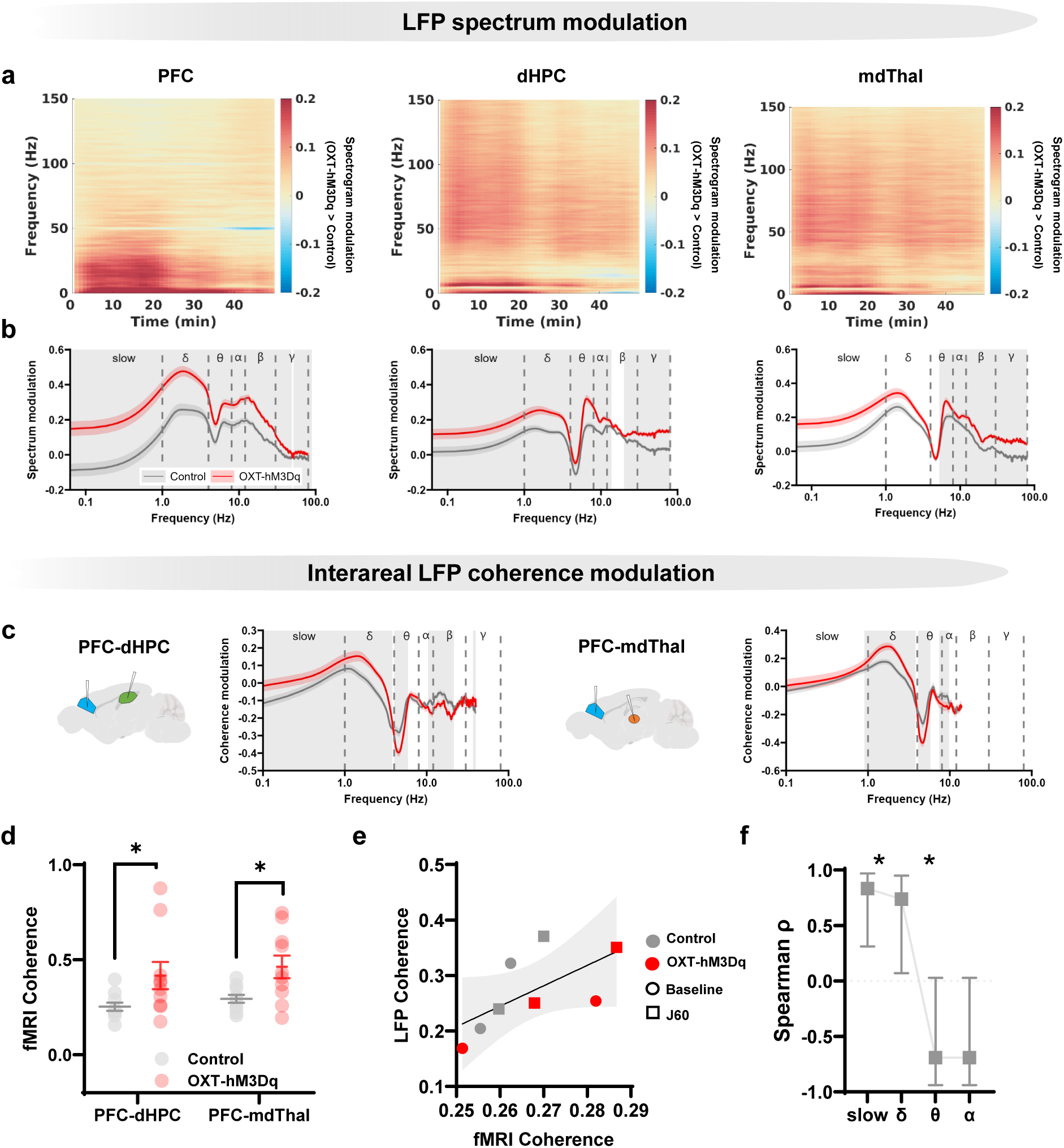
Endogenous oxytocin modulates local electrophysiological activity and interareal synchrony. (a) Differences in time resolved LFP spectrogram modulation between groups (OXT-hM3Dq – Control, J60 administration at time 0). (b) Frequency-resolved LFP spectrogram modulation after J60 administration. Gray shading indicates frequency clusters surviving FWER correction (Wilcoxon rank-sum tests, grey bands indicate FWER-corrected clusters with p < 0.001). (c) Frequency-resolved interareal LFP coherence modulation between mPFC–HPC and PFC– Thal following J60 administration. Gray shading indicates frequency clusters surviving FWER correction (Wilcoxon rank-sum tests, grey bands indicate FWER-corrected clusters p < 0.001). (d) Low-frequency BOLD fMRI coherence (0.01– 0.015 Hz) between PFC and hippocampus or thalamus after J60 administration. Individual animals and group mean ± 95% CI are shown; paired t-test, *p < 0.005. (e) Correlation (Spearman) between low-frequency (0.01 - 0.1Hz) LFP coherence and BOLD signal coherence. Dots indicate group averages at baseline (i.e., before J60, circle) and after j60 (square) for matched region-electrode pairs. Linear fit ± 95% CI is shown. (f) Band-resolved Spearman correlation between LFP coherence BOLD signal coherence (*p < 0.05). PFC, prefrontal cortex; Thal, thalamus; dHPC, dorsal hippocampus.

We next examined whether OXT-dependent changes in local electrophysiological activity were accompanied by altered interareal coupling. To this end, we computed fronto-hippocampal and fronto-thalamic LFP coherence over frequencies exceeding the coherence noise floor. We found that endogenous OXT release elicited bidirectional spectral changes in interareal LFP coherence in both region pairs, with increased coherence at low frequencies and reduced coherence at higher frequencies (Wilcoxon rank-sum tests, cluster-corrected p < 0.001; Figure 4c). Recent work has shown that slow electrophysiological coherence scales positively with resting-state fMRI connectivity (Rocchi et al., 2022; Sastre-Yagüe et al., 2026; Wang et al., 2012). We thus asked whether OXT-dependent changes in low-frequency electrophysiological coupling could account for the observed fMRI connectivity changes. Direct comparisons of LFP coherence and fMRI connectivity, however, revealed an apparent mismatch: low-frequency LFP coherence increased in both region pairs, whereas Pearson-based fMRI connectivity increased for PFC-thalamus, but became negative for PFC-hippocampus. We reasoned that this apparent discrepancy may reflect the different nature of the two measures. Coherence magnitude captures frequency-specific synchrony irrespective of whether signals are in phase or anti-phase, whereas Pearson correlation-based fMRI connectivity is a signed zero-lag measure of covariance, and as such it is sensitive to temporal alignment between signals.

We therefore asked whether fMRI coherence, rather than zero-lag Pearson correlation, would better match the observed electrophysiological effects. Consistent with this hypothesis, fMRI coherence increased in both PFC-thalamus and PFC-hippocampus pairs following endogenous OXT release (paired t-test in the 0.01–0.015 Hz range, p < 0.005; Figure 4d). Notably, when we examined the relationship between fMRI and LFP coherence across frequency bands and region pairs (Figure 4e), we found significant positive associations between these two measures in the slow and δ bands, but not at higher frequencies (Figure 4f). Together, these findings indicate that OXT-dependent fMRI connectivity changes are associated with corresponding changes in low-frequency electrophysiological coherence, albeit with different phase relationships across region pairs.

### Endogenous OXT biases fMRI state dynamics

Our analyses so far show that endogenous OXT release reconfigures functional connectivity across distributed brain networks. We next asked whether these connectivity changes reflect a static reorganization of the “fMRI connectome”, or instead emerge from altered dynamics of recurrent fMRI activity states. To this end, we analyzed the dynamic organization of fMRI activity by decomposing it into recurring co-activation modes (c-Modes)(Gutierrez-Barragan et al., 2022) (Figure 5a). This approach allowed us to parsimoniously describe fMRI network dynamics as transitions among five dominant recurring fMRI states, which together explained more than 60% of the variance in the fMRI time series (see Methods).

**Figure 5.**
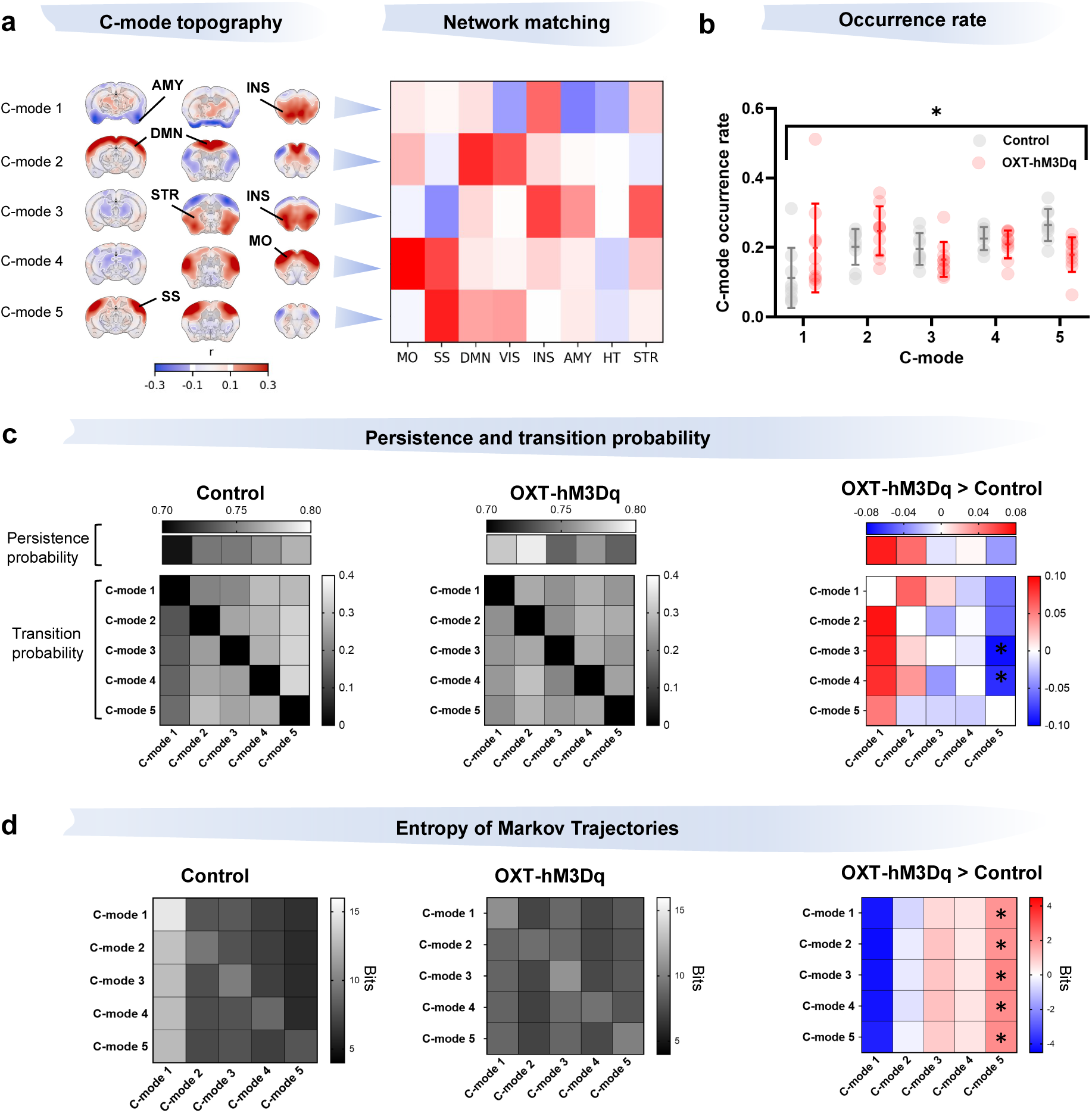
Endogenous oxytocin reorganizes large-scale brain network dynamics. (a) C-mode topographies (left) and corresponding network coactivation profiles (right). Network profiles were obtained by extracting mean C-mode values within predefined network masks and spatially z-scoring the resulting values. (b) C-mode occurrence rates in control and OXT-hM3Dq mice. Data are shown as mean ± 95% CI. Group differences in C-mode occurrence were assessed using a MANOVA on isometric-log ratio transformed occurrence rates (*p < 0.05). (c) C-mode persistence probabilities (top rows) and transition (off-diagonal) probabilities (matrices) between distinct C-modes in control and OXT-hM3Dq mice. Right, corresponding between-group differences (OXT-hM3Dq − control). *q < 0.05, FDR-corrected permutation test. (d) Entropy of Markov trajectories (HMT) between C-mode states and between-group differences. Higher entropy indicates lower accessibility of a destination C-mode from a given starting C-mode. *FDR < 0.05, permutation test.MO, motor; SS, somatosensory; DMN, default mode network; VIS, visual network; INS, insula; AMY, amygdala; HT, hypothalamus; STR, striatum.

We first assessed whether endogenous OXT release altered the temporal organization of these C-modes. J60 administration elicited a significant redistribution of C-mode occurrence rates in OXT-hM3Dq mice relative to controls (MANOVA on isometric-log ratio transformed occurrence rates, p < 0.05; Figure 5b). This shift was marked by higher occurrence of C-mode 1, a state with prominent salience-related anterior insular components, and reduced occurrence of C-mode 5, a somatosensory-dominated state. Notably, OXT release also altered the structure of state transitions, with increased persistence probability of C-mode 2 (t-test, permutation p<0.05; Figure 5c), indicating enhanced stability of this DMN-like state, and increased bidirectional transition probabilities between C-mode 2 and salience-related C-mode 1 (t-test, permutation p<0.05; Figure 5c). Conversely, OXT release reduced transitions from C-mode 3 and C-mode 4 to somatosensory-dominated C-mode 5 (t-test, permutation p < 0.05; Figure 5c). This transition architecture was associated with changes in C-mode accessibility as tracked by entropy of corresponding Markov trajectories. Here, lower entropy values indicate more direct and deterministic transition paths to a target state, whereas higher entropy values indicate more uncertain trajectories involving intermediate C-modes. Consistent with occurrence-rate results, endogenous OXT release reduced entropy for transitions terminating in C-mode 1, indicating increased accessibility of this salience-related state, and increased trajectory entropy for transitions terminating in C-mode 5 (p < 0.05; Figure 5d), indicating reduced accessibility of this somatosensory-dominated state. Together, these findings indicate that endogenous OXT release biases fMRI brain-state dynamics by increasing engagement of salience-related states, and by concomitantly reducing the accessibility of somatosensory-dominated states.

## Discussion

In the present study, we combined cell-type-specific chemogenetics, fMRI and *in vivo* electrophysiology to probe how endogenous OXT-neuron activation shapes brain-wide functional organization. We found that endogenous OXT release produces widespread hemodynamic activation, together with a focal reorganization of large-scale functional connectivity marked by increased occurrence of salience-related states. These network effects were accompanied by robust electrophysiological changes, including increased low-frequency interregional coupling mirroring corresponding fMRI coherence changes. Together, these findings identify functional connectivity reconfiguration and dynamic network-state biasing as a systems-level mechanism through which OXT can tune large-scale socio-affective circuits.

Much of what we know about the brain-wide effects of OXT comes from human and animal neuroimaging studies based on exogenous administration of the peptide, most commonly via intranasal delivery (Galbusera et al., 2017; Quintana et al., 2021). These studies have been instrumental in establishing that OXT can, in principle, modulate large-scale brain activity and functional connectivity. Perfusion MRI has revealed distributed and temporally structured changes in regional cerebral blood flow across limbic, striatal and frontocortical territories following intranasal OXT administration (Martins et al., 2020; Paloyelis et al., 2016). Resting-state fMRI studies further revealed that intranasal OXT can produce network-specific and bidirectional changes in functional connectivity, encompassing cohort-dependent increases in amygdala–prefrontal or amygdala–cingulate coupling (Ebner et al., 2016; Kovacs & Keri, 2015), along with altered integration and segregation across salience, default-mode and attention networks, and regionally selective changes in functional connectome organization (Brodmann et al., 2017; Ebner et al., 2016; Galbusera et al., 2017; Kovacs & Keri, 2015; Liu et al., 2022; Martins et al., 2021; Pagani et al., 2020; Wu et al., 2023; Xin et al., 2021). However, exogenous administration does not directly isolate the consequences of activating endogenous OXT-releasing neurons, and the ensuing functional effects can depend on dose, route of administration, and variability in central engagement following intranasal delivery (Quintana et al., 2021). As a result, the systems-level consequences of endogenous OXT-neuron activation have remained largely unexplored.

By leveraging cell-type-specific stimulation of OXT-producing neurons in mice, our study provides an experimental framework that enables the identification of the whole-brain substrates engaged by *endogenous* OXT signaling in the mammalian brain. This distinction is key, because endogenous OXT-neuron activation and exogenous OXT administration may differ not only in the way they engage the OXT system, but also in the kinetics and magnitude of the resulting brain response. For example, previous rCBV mapping in mice showed that intranasal OXT elicited a biphasic hemodynamic response characterized by an early, spatially extended activation followed by a sustained, more focal response encompassing basal forebrain and hippocampal regions (Galbusera et al., 2017). In the present study, chemogenetically evoked OXT-neuron activation instead produced a sustained and widespread hemodynamic response across many subcortical and cortical areas. While this difference may partly reflect the distinct temporal profiles of chemogenetic stimulation and intranasal delivery, these results nonetheless indicate that the two paradigms may engage brain-wide activity with markedly different kinetics, and spatial distribution.

The widespread rCBV response elicited by endogenous OXT-neuron activation was not readily predicted by OXT projection density or OXTR expression, arguing against a point-to-point model in which OXT acts only at directly innervated, receptor-rich targets. This interpretation is consistent with anatomical evidence showing a quantitative mismatch between OXT projections and OXTR expression across the mouse brain (Son et al., 2022). Thus, OXT-evoked brain-wide activation may reflect the combined contribution of local axonal release, volume- or CSF-mediated OXT signaling, and indirect recruitment of polysynaptic pathways activated by OXT itself, or by co-released transmitters, such as glutamate (Hasan et al., 2019). Irrespective of the precise mechanism, our findings suggest that endogenous OXT-neuron activation can recruit distributed brain systems in a manner not reducible to anatomical projection density or OXTR alone.

Importantly, the widespread hemodynamic response produced by endogenous OTX was accompanied by a much more selective reorganization of fMRI connectivity across hypothalamic, amygdalar, prefrontal, and hippocampal regions. This dissociation suggests that broad OXT-driven regional recruitment can be translated into more selective changes in interareal communication, providing a potential network-level mechanism through which OXT engages specific functional circuits. Consistent with this interpretation, previous work has directly implicated these regions in OXT-dependent social and affective functions. For example, OXT signaling in hypothalamic nuclei has been shown to contribute to social recognition memory (Thirtamara Rajamani et al., 2024), while OXT action in hippocampal, amygdalar, and prefrontal circuits has been linked to social memory encoding (Tsai et al., 2022), affective and social processing (Ferretti et al., 2019; Knobloch et al., 2012), and social recognition (Tan et al., 2019), respectively. Our imaging results thus suggest that endogenous OXT can selectively reshape interareal coupling among regions implicated in these social-affective functions.

Notably, these fMRI effects were accompanied by pronounced changes in electrophysiological activity, arguing against a purely vascular origin for the observed connectivity changes. Specifically, endogenous OXT altered local LFP power and reorganized interareal electrophysiological coupling, producing a robust increase in low-frequency coherence across fronto-thalamic and fronto-hippocampal circuits. These effects were accompanied by heterogeneous changes in multi-unit activity, indicating that OXT-neuron activation altered the temporal organization of both local and distributed neural activity, without uniformly scaling population firing. This interpretation is consistent with previous *ex vivo* electrophysiological work showing that OXT can reshape local circuit dynamics by modulating spike timing, transmission fidelity, and burst firing (Owen et al., 2013; Tirko et al., 2018). In keeping with recent work linking slow electrophysiological synchrony to resting-state fMRdI connectivity (Rocchi et al., 2022; Sastre-Yagüe et al., 2026), we also found that low-frequency LFP coherence covaried positively with fMRI coherence across region pairs. This cross-modal correspondence provides a possible electrophysiological correlate of the observed OXT-dependent fMRI connectivity changes. Interestingly, this correspondence did not extend to zero-lag fMRI connectivity, as prefrontal-hippocampal fMRI connectivity (i.e. Pearson correlation) became negative upon endogenous OXT release, whereas LFP and fMRI coherence between these two regions increased. This dissociation suggests that negative fMRI connectivity in this circuit may reflect altered temporal alignment within coordinated activity, rather than functional decoupling. A contribution from region-specific neurovascular coupling should also be considered, as hippocampal neurovascular responses are weaker and less tightly coupled to neural activity than neocortical responses in awake mice (Shaw et al., 2021). Negative hippocampal BOLD responses have also been reported despite increased neuronal and hemodynamic activity (Schridde et al., 2008), further highlighting the complexity of neurovascular relationships in this region. While the precise origin of the observed anticorrelation remains unresolved, our observations caution against interpreting negative fMRI connectivity as simple functional decoupling.

Importantly, our dynamic fMRI analyses extend this multiscale account by showing that endogenous OXT release robustly alters the temporal organization of recurrent fMRI brain states as probed within the C-mode framework (Gutierrez-Barragan et al., 2024). Specifically, OXT increased the occurrence of salience-related states while reducing that of somatosensory-dominated states. Trajectory-entropy analyses revealed that this was paired with increased and decreased accessibility of salience and somatosensory-related states, respectively. Together, these findings suggest that endogenous OXT-neuron activation biases the functional state-space of the brain, potentially tuning the brain’s responsiveness to salient inputs. This interpretation is broadly consistent with human EEG microstate studies showing that intranasal OXT increases the representation of microstates associated with attention, while reducing configurations linked to internally oriented processing (Schiller et al., 2019; Zelenina et al., 2022).

Social-salience and allostatic accounts propose that OXT does not impose a fixed prosocial state, but adjusts information processing and behavioral readiness according to internal state and environmental demands (J. A. Bartz et al., 2011; Menon & Neumann, 2023; Quintana & Guastella, 2020; Shamay-Tsoory & Abu-Akel, 2016). Our findings suggest a potential network-level basis for this modulation. Within this framework, OXT could flexibly tune behavioral responses to contextually relevant inputs by changing which recurrent brain states are preferentially expressed and accessed, without simply activating a fixed set of target regions. Future studies combining endogenous OXT-neuron manipulations with imaging or electrophysiology during behavior will be needed to determine whether and how this state-space reweighting translates into social, affective, or motivational behavior.

In conclusion, our findings show that endogenous OXT release produces widespread rCBV activation, selectively reorganizes functional connectivity, increases low-frequency interregional synchrony, and biases recurrent fMRI dynamics toward more accessible salience-related states. These results support a model in which OXT may tune behavior not merely by modulating regional activity, but by reshaping interregional coupling and the accessibility of distributed network configurations.

## Acknowledgments

This work has been funded by the European Research Council (ERC) under the European Union’s Horizon 2020 research and innovation program (no. 101125054 #BRAINAMICS, no. 802371 #DISCONN to A.Gozzi), and by the Brain and Machines Flagship Program of the Italian Institute of Technology. E.d.G. acknowledges funding from the Canadian Institutes of Health Research (CHIR, MFE-187902).

## Materials and Methods

### Ethical statement

All animal procedures were conducted in accordance with the Italian Legislative Decree 26/2014 and EU Directive 2010/63/EU, and followed the recommendations of the Guide for the Care and Use of Laboratory Animals. Experimental protocols were reviewed and approved by the Animal Care Committee of the Istituto Italiano di Tecnologia and by the Italian Ministry of Health. All surgical procedures were performed under anesthesia. Mice were housed under controlled temperature (21 ± 1°C) and humidity (60 ± 10%) conditions, with food and water available *ad libitum*.

### Mouse Line Generation

To enable selective chemogenetic stimulation of oxytocin (OXT)-releasing neurons, mice harboring a conditional ROSA26-CAG-DIO-hM3Dq-mCherry allele (Giorgi et al., 2017) were crossed with OXT-Cre mice, in which Cre recombinase is inserted downstream of the endogenous oxytocin locus (Wu et al., 2012). In the resulting OXT-hM3Dq mice, Cre-mediated recombination induces the expression of the excitatory DREADD receptor hM3Dq selectively in OXT-producing neurons, enabling their activation following administration of a DREADD ligand.

The OXT-hM3Dq line was maintained by crossing hM3Dq-positive, Cre-negative mice with hM3Dq-negative, OXT-Cre-positive mice. This breeding strategy yields OXT-hM3Dq mice heterozygous for the hM3Dq allele and corresponding littermate controls. Mice were routinely genotyped by PCR DNA amplification with specific oligonucleotides as primers. For the ROSA26 wild-type allele, the forward primer was 5′-GAGGGGAGTGTTGCAATACC-3′ and the reverse primer was 5′-AGTCTAACTCGCGACACTGTA-3′. The DIO-hM3Dq allele was detected using the same forward primer and the alternative reverse primer 5′-GTCCCTATTGGCGTTACTATG-3′. For the OXT-Cre locus, the forward primer was 5′-TTTGCAGCTCAGAACACTGAC-3′ and the wild-type reverse primer was 5′-AGCCTGCTGGACTGTTTTTG-3′. Cre insertion downstream of the Oxt stop codon was detected using the same forward primer and the alternative reverse primer 5′-ACACCGGCCTTATTCCAAG-3′. Adult male OXT-hM3Dq mice positive for both the hM3Dq allele and OXT-Cre transgene were used as the experimental group, whereas littermate male mice carrying either transgene alone or neither transgene served as controls. Experiments were performed in male mice, except for ex vivo patch-clamp recordings, which included both sexes.

### Immunohistochemistry on OXT-Rosa26-tdTomato reporter line

In a separate validation cohort, we crossed Oxt-IRES-Cre mice with the Rosa26-tdTomato reporter line and performed immunohistochemistry to assess the overlap between Cre-dependent tdTomato expression and oxytocin-immunoreactive neurons (Figure 1a). Male mice from this cohort were perfused transcardially with 4% PFA, and 50 μm -thick brain coronal sections were obtained with a vibratome. Free-floating sections were incubated overnight at 4 °C with a primary rabbit anti-RFP antibody (ab62341, Abcam, 1:1000) and with a primary mouse anti-Oxt antibody (MAB5296, Millipore, 1:1000). Sections were then incubated overnight at 4 °C with a mouse Alexa Fluor 488 antibody (O-11033, ThermoFisher 1:500) and a rabbit Rhodamine Red antibody (R6394, ThermoFisher 1:500).

### Immunohistochemistry and in situ hybridization on OXT-hM3Dq mice

For immunohistochemistry and *in situ* hybridization analyses adult male OCT-hM3Dq mice (n = 5) were perfused transcardially with 4% paraformaldehyde (PFA). Brains were dissected and post-fixed overnight at 4 °C. Brains were then equilibrated in 30% sucrose and sectioned at 18 µm using a cryostat. Parallel coronal sections were used for immunohistochemistry and in situ hybridization. For immunohistochemistry, sections were incubated overnight at 4°C with rabbit anti-RFP antibody (ab62341, Abcam, 1:1000), followed by overnight incubation at 4°C with Rhodamine Red-conjugated anti-rabbit secondary antibody (R6394, ThermoFisher, 1:500). *In situ* hybridization was performed as previously described (Migliarini et al., 2013), using a digoxigenin-labeled Oxt antisense riboprobe (0.4 kb). Signal was developed using NBT/BCIP substrate for alkaline phosphatase (11681451001, Roche). Cell counts were obtained from five animals, encompassing 34 sections processed with each method, for a total of 68 sections. Data are expressed as the percentage of RFP-positive cells relative to Oxt-positive cells. Equivalence between RFP-positive and Oxt-positive cell counts was assessed using a mixed-effects two one-sided tests (TOST) procedure with predefined equivalence bounds of ±10%.

### Drug formulation and experimental groups

The water-soluble DREADD ligand JHU37160 dihydrochloride (J60, HB6261, Hello Bio, (Bonaventura et al., 2019)), was dissolved in sterile saline at a concentration of 0.3 mg/ml. J60 was administered intraperitoneally at a dose of 1 mg/kg (injection volume: 3,3 µl/g body weight). Both wild-type and OXT-hM3Dq mice received JHU37160 treatment to account for potential off-target effects of the DREADD agonist. J60 instead was used instead of clozapine-N-oxide (CNO) because peripherally administered CNO can be reverse-metabolized to clozapine in vivo (Gomez et al., 2017), and clozapine has previously been shown to elicit peripheral release of OXT (Uvnas-Moberg et al., 1992).

### Ex vivo patch-clamp electrophysiology

Mice were anesthetized with isoflurane and transcardially perfused with ice-cold cutting solution containing: 200 mM sucrose, 4 mM MgCl_2_, 2.5 mM KCl, 1.25 mM NaH_2_PO_4_, 0.5 mM CaCl_2_, 25 mM NaHCO_3_ and 10 mM D-glucose (∼300 mOsm, pH 7.4, oxygenated with 95% O_2_ and 5% CO_2_). Brains were removed and immersed in cutting solution. Coronal slices (270 um cut with VT1000S Leica Microsystem vibratome) were allowed to recover for 1 hour at 35°C in a solution containing: 117 mM NaCl, 2.5 mM KCl, 1.25 mM NaH_2_PO_4_, 3 mM MgCl_2_, 0.5 mM CaCl_2_, 25 mM NaHCO_3_ and 10 mM glucose (∼310 mOsm, pH 7.4, oxygenated with 95% O_2_ and 5% CO_2_).

Recordings were performed in visually identified neurons of the PVN at room temperature in artificial cerebrospinal fluid (ACSF), with the following composition: 117 mM NaCl, 2.5 mM KCl, 1.25 mM NaH_2_PO_4_, 1 mM MgCl_2_, 2 mM CaCl_2_, 25 mM NaHCO_3_ and 10 mM glucose (∼310 mOsm, pH 7.4, oxygenated with 95% O_2_ and 5% CO_2_). Patch pipettes were made from thick-wall borosilicate glass capillaries (B150-86-7.5, Sutter Instrument). Pipettes (5-7 MΩ) were filled with intracellular solution containing: 130 mM K-gluconate, 10 mM HEPES, 7 mM KCl, 0.6 mM EGTA, 4 mM Mg_2_ATP, 0.3 Mm Na_3_GTP, 10 mM Phosphocreatine. The pH was adjusted to 7.3 with HCl. Once stable recording conditions were obtained, with series resistance <20 MΩ, PVN neurons were classified electrophysiologically as magnocellular or parvocellular based on the presence or absence of transient outward rectification, respectively (Supplementary Figure 2), using an established current-clamp protocol (Ferretti et al., 2019; Luther et al., 2000). DREADD activation was induced by bath application of 10 μM clozapine-N-oxide (CNO; #4936, Tocris Bioscience) for 10 min. CNO was used only for *ex vivo* validation of hM3Dq function in acute slices, where reverse metabolism to clozapine is not relevant (Gomez et al., 2017). Frequency discharge was calculated as firing frequency (Hz) after 5 minutes of CNO application. Data were acquired with a patch-clamp amplifier (Multiclamp 700B, Molecular Devices) filtered at 0.1 Hz and 5 kHz, sampled at 10 kHz, and analyzed using pClamp 10.2 software (Molecular Devices). All chemicals were purchased from Sigma, unless otherwise specified.

We analyzed n = 12 cells from 5 wild-type mice (3 females and 2 males) and n= 37 cells from 6 OXT-hM3Dq mice (4 females and 2 males). As OXT-expressing neurons comprise only a subset of cells within the PVN, and mCherry fluorescence could not be visualized under the recording conditions due to low fluorescence of the fusion protein, transgene-expressing (hM3Dq+) cells could not be identified prospectively. We therefore sampled PVN cells without prior selection, with the expectation that a proportion would not express hM3Dq and consequently would not respond to CNO. We first evaluated the effect of CNO across all recorded PVN cells, without selection based on their response, and observed a significant increase in firing frequency (Supplementary Figure 3). To characterize the CNO responsive subgroup, we included the 18 cells that showed an increase in frequency discharge >0.5 Hz after CNO application (Figure 1c).

### Plasma OXT quantification

Plasma oxytocin levels were quantified by radioimmunoassay as previously described (Neumann et al., 2013). Briefly, the left femoral artery was cannulated under isoflurane anesthesia. After surgery, isoflurane was replaced with 0.8% halothane to match the sedation protocol used for fMRI experiments. Thirty minutes after the isoflurane-to-halothane switch, a baseline arterial blood sample (0.15 ml) was collected through the femoral artery from male control (n = 5) and OXT-hM3Dq (n = 5) mice. Blood withdrawal was followed by fluid volume replacement with sterile saline. Five minutes later, JHU37160 (1 mg/kg, i.p.) was administered, and a second arterial blood sample (0.15 ml) was collected 40 min post-injection.

Blood samples were centrifuged (10 min, 1300 g, 4°C) and plasma samples (approximately 0.1 ml) were stored at -20°C until the radioimmunoassay procedure was performed. OXT levels were estimated in plasma samples using a radioimmunoassay (RIAgnosis, Munich) as previously described (Galbusera et al., 2017; Neumann et al., 2013).

### Contrast-enhanced fMRI

Animal preparation for contrast-enhanced fMRI has been previously described in detail (Ferrari et al., 2012; Squillace et al., 2014). Experiments were carried out in n = 10 control and n = 11 OXT-hM3Dq male mice under light halothane sedation. We employed halothane-based light sedation as this procedure ensures stable brain states over long recordings, preserves cerebral autoregulation, as well as resting-state fMRI network organization in mice, with good cross-species correspondence with those observed in human imaging studies (Alvino et al., 2025; Bertero et al., 2018; Coletta et al., 2020; Gozzi et al., 2025).

Animals were anaesthetized with isoflurane (5% induction), intubated, and artificially ventilated (2%). The tail vein was cannulated for intravenous administration of the contrast agent. An intraperitoneal catheter was inserted during animal preparation for J60 administration. At the end of preparation, isoflurane was discontinued and replaced with halothane (0.8%). Functional data acquisition started 30 min after isoflurane cessation. We used a 7 T MRI scanner (Bruker, Ettlingen) equipped with a BGA-9 gradient set (380 mT/m, max. linear slew rate 3,420 T/m/s), a 72 mm birdcage transmit coil, and a 4-channel solenoid receive coil. The scanner was operated using Paravision 6.01 software (Bruker, Ettlingen). fMRI images were acquired using a Fast Low-Angle Shot (FLASH) sequence (TReff = 395 ms, TEeff = 3 ms, α = 30°; matrix size, 192 × 192 × 24, resolution, 0.156 x 0.156 x 0.5 mm, 4 averages, dt = 60 s, N = 90 corresponding to 90 min total acquisition time). To sensitize the fMRI signal to changes in relative cerebral blood volume (rCBV), mice received an intravenous injection of superparamagnetic iron oxide nanoparticles (Molday ION, BioPAL, Worcester, MA, USA; 5 μL/g body weight), as previously described (Galbusera et al., 2017; Squillace et al., 2014). The injection of the contrast agent and J60 (1 mg/kg, i.p.) was performed 5 and 30 minutes after scan start, respectively.

Changes in rCBV were quantified as previously described (Errico et al., 2015; Galbusera et al., 2017; Giorgi et al., 2017). In particular, fMRI time series were spatially normalized to a common reference space, and signal intensity was converted to fractional rCBV changes relative to the 8-min baseline preceding J60 administration, without detrending. Voxel-wise group statistics were performed using a step function depicting a transition at 22 min following J60 administration, corresponding to the onset of the average rCBV response observed in OXT-hM3Dq mice. Group comparisons between OXT-hM3Dq and control mice were performed using multilevel Bayesian inference with a cluster forming threshold of Z > 2.1 and a cluster significance threshold of p < 0.05 (FEAT Version 6.00).

### Resting-state fMRI

Resting-state BOLD fMRI scans were acquired from male control mice (n = 10) and OXT-hM3Dq mice (n = 10) using the same 7 T scanner and sedation/ventilation protocol described above for contrast-enhanced rCBV fMRI. An intraperitoneal catheter was inserted during animal preparation for J60 administration. Resting-state fMRI time series were acquired using a single-shot echo-planar imaging sequence with the following parameters: TR/TE = 1000/15 ms, flip angle = 60°, matrix size = 98 × 98, field of view = 2.3 × 2.3 cm², 18 coronal slices, and slice thickness = 550 μm, slice gap = 50um. A total of 4,620 volumes were acquired over 77 min, including 17 min before and 60 min after intraperitoneal J60 administration.

### fMRI connectivity mapping

Resting-state fMRI data were preprocessed as previously described (Gutierrez-Barragan et al., 2024) using a combination of AFNI (Cox & Hyde, 1997), ANTs (Tustison et al., 2021) and FSL (Jenkinson et al., 2012) tools. The first 120 volumes (2 min) were removed to allow for signal equilibration. Data were then despiked (AFNI 3dDespike), motion-corrected (FSL MCFLIRT), and registered to an in-house mouse brain template (0.23 × 0.23 × 0.6 mm³ resolution; ANTs registration suite). Nuisance regression was performed using AFNI 3dDeconvolve and included the mean cerebrospinal fluid signal, together with 24 motion regressors (i.e., the six rigid-body motion parameters, their temporal derivatives, and the squared terms of both the original parameters and their derivatives). Time series were band-pass filtered (0.01–0.1 Hz; AFNI 3dBandpass), spatially smoothed (0.5 mm FWHM kernel; AFNI 3dBlurInMask), and each voxel time series was normalized to z-scores. Quantifications of between-group differences in fMRI connectivity were carried out on the 15 and 60 min window pre and post J60 injection, respectively.

To obtain a spatially unbiased identification of the brain regions exhibiting alterations in functional connectivity, we calculated voxelwise global functional connectivity maps as the mean Fisher-z-transformed temporal correlation between a given voxel and all other brain voxels. Correlation coefficients were Fisher-transformed prior to averaging. Group differences were assessed voxel-wise using two-sample tests, and statistical maps were thresholded at |Z| > 2.1, cluster-corrected p < 0.05). Interregional functional coupling was further assessed by computing region-wise connectivity matrices based on a previously defined mouse brain parcellation (Gutierrez-Barragan et a., 2022). Pearson correlation coefficients were calculated between all pairs of regions, and Fisher transformed. Regions were assigned to previously described functional networks and modules, including the default-mode, hippocampal, latero-cortical, and olfactory-basal forebrain networks (Coletta et al., 2020). Seed-based functional connectivity analyses were performed using regions of interest centered on the hypothalamus, prefrontal cortex, and dorsal hippocampus. For each seed region, voxel-wise Pearson correlation maps were computed and Fisher-transformed prior to statistical testing. Group differences were assessed using voxel-wise two-sample t-tests (|Z| > 2.1, cluster-corrected p < 0.05).

### In vivo electrophysiological recordings

*In vivo* electrophysiological recordings were acquired in male OXT-hM3Dq (n = 9) and control (n = 11) mice under the same sedation/ventilation protocol used for rsfMRI. Mice were anesthetized with isoflurane (5% induction), intubated, artificially ventilated (2%), and head-fixed in a stereotaxic frame (Stoelting). Craniotomies were performed above the right prefrontal cortex (PFC; AP +1.8 mm, ML +0.35 mm) and dorsal hippocampus (AP −1.8 mm, ML +1.8 mm). A single-shank silicon probe (Cambridge Neurotech H7b, 32 channels, 25 µm contact spacing) was lowered vertically into the medial PFC (depth 1.9 mm). A second silicon probe (Cambridge Neurotech L3, 64 channels, 50 µm contact spacing) was inserted at a 60° angle to a depth of 4 mm targeting the dorsal hippocampus and mediodorsal thalamus. Before insertion, probes were coated in DiI (thermofisher). Probes were advanced using a motorized microdrive system (New Scale Technologies) at 2-4 µm/s. Ground electrodes were placed in contact with cerebrospinal fluid through a craniotomy above the cerebellum.

During electrode insertion, isoflurane was discontinued and replaced with halothane sedation (0.8%). Electrophysiological data acquisition began 1.5 h after isoflurane cessation, corresponding to 1 h after completion of electrode insertion. This interval was used to allow isoflurane washout and to minimize residual burst-suppression activity associated with prolonged exposure to the deeper anesthesia required for electrode insertion. After 30 minutes of baseline recording, J60 was injected intraperitoneally, and recordings continued for 60 minutes after J60 administration. Signals were amplified using an RHD 2000 amplifier system (Intan Technologies, RHD Recording Controller Software, v2.09) and sampled at 30 kHz.

Following completion of the recordings, mice were deeply anesthetized with pentobarbital and transcardially perfused with phosphate-buffered saline (PBS), followed by 4% methanol-free formaldehyde. Brains were dissected and post-fixed overnight at 4 °C in the same fixative. Coronal brain sections encompassing both probe insertion sites were cut at 50 μm thickness using a vibratome and counterstained with DAPI. Probe trajectories were verified by fluorescence microscopy based on DiI staining and reconstructed using the Neuropixels Trajectory Explorer (DOI: 10.5281/zenodo.7043459). Using the probe length, insertion point, insertion angle, and insertion depth, the anatomical location of each recording site (“Probe area” in the Trajectory Explorer) was assigned according to the Allen Common Coordinate Framework. Region assignments were exported separately for each mouse to account for minor variability in probe placement and used for all subsequent analyses.

### Local field potential and multi-unit activity

Local field potentials (LFP) and multi-unit activity (MUA) were processed as previously described (Belitski et al., 2008; Rocchi et al., 2022; Sastre-Yagüe et al., 2026). Signals were filtered in the forward and reverse directions to avoid phase distortion and resampled to 1,000 and 5,000 Hz for LFP and MUA analyses, respectively. Spike times were detected from the high-frequency signal using a threshold corresponding to four times the median of the absolute signal divided by 0.6745 (Quiroga et al., 2004). Events occurring within 1 ms of a preceding spike were excluded.

Multi-unit firing rates were computed for each recording channel in 1-min epochs, spanning 20 min of stable baseline, and a 50 min time window post J60 administration. The first 5 min following J60 administration were excluded to remove injection-related motion artifacts, and the final 5 min of the recording were discarded to harmonize recording duration across animals. For statistical analyses, firing rates were log-transformed and then baseline-normalized by dividing each value by the mean log-transformed baseline firing rate of the corresponding channel. Average firing rates were computed for the pre- and post-injection periods. Group differences across periods were analyzed using a linear mixed-effects model with fixed effects of group, time, and their interaction, and random intercepts for channels nested within animals (firing rate ∼ group × time + (1|animal/channel)). For visualization, firing rates across time were averaged across channels and animals within each group and displayed with bootstrapped confidence intervals.

LFP power spectra were computed in consecutive 1-min epochs. For each epoch, power spectra were calculated separately for each channel and then averaged across channels within a given anatomical region, yielding one spectrum per subject, region and epoch. Baseline spectra were obtained by averaging all pre-injection epochs. Spectral modulation following J60 administration was quantified for each post-injection epoch using a modulation index defined as (Ppost − Pbaseline)/(Ppost + Pbaseline), where Ppost and Pbaseline denote the power spectra of the post-J60 and baseline periods, respectively. At each frequency, spectral modulation values for each epoch from control and OXT-hM3Dq mice were compared using a two-sample t-test. The resulting t-statistics were thresholded at α = 0.05 and contiguous suprathreshold frequency bins were grouped into clusters. For each cluster, a cluster-mass statistic was calculated as the sum of t-values within the cluster. Statistical significance was assessed using a permutation test in which group labels were randomly shuffled, generating a null distribution of cluster-mass values against which the observed clusters were evaluated (Maris & Oostenveld, 2007).

Magnitude-squared coherence was calculated between all pairs of electrode channels using Welch’s averaged periodogram method (mscohere, MATLAB). Coherence spectra were estimated in consecutive 1-min epochs for all channel pairs using 2000-sample windows with 1000-sample overlap. For each subject, region pair and epoch, coherence spectra were averaged across all channel pairs connecting the two regions. The baseline coherence spectrum was obtained by averaging the coherence of all pre-injection epochs. Coherence modulation following J60 administration was then quantified for each post-injection epoch using a modulation index calculated as (Cpost − Cbaseline)/(Cpost + Cbaseline), where Cpost and Cbaseline denote the coherence spectra during the post-J60 and baseline periods, respectively. Statistically significant differences between groups were assessed using the frequency-resolved cluster-based correction approach described above for LFP spectrum modulation.

Coherence analyses were restricted to region pairs and frequency ranges exhibiting significant baseline coherence. Significance thresholds were estimated using surrogate data generated through circular time shifts. To estimate the noise floor distribution for each frequency, coherence was computed for 25 random channel pairs, each with 200 random circular time shifts, for a total of 5,000 shuffled time series. For each frequency, the noise-floor threshold was defined as the average, across epochs, of the 95th percentile of the surrogate coherence distribution. Based on this, analyses were restricted to frequencies below 40 Hz for PFC-hippocampal coupling, and below 13 Hz for PFC-mediodorsal thalamic coupling.

To compare electrophysiological and fMRI measures of interareal coupling, coherence of BOLD fMRI data was computed between region pairs corresponding to the electrophysiology recording locations (PFC-HPC, and PFC-Thal, respectively). fMRI coherence was calculated separately for the baseline and post-J60 periods. For the between-group comparison (Figure 4d), coherence was averaged over 0.01–0.015 Hz, i.e., the frequency range in which between-group differences were observed in a frequency-resolved analysis. For correlations with electrophysiology (Figure 4e,f), coherence was averaged over the full analyzed BOLD frequency range (0.01–0.1 Hz) and then within each group for each region pair and time period. Electrophysiological coherence was quantified as described above and averaged separately for the baseline and post-J60 periods for each group, region pair and frequency band: slow 0.1-1 Hz, δ 1-4 Hz, θ 4-8 Hz, α 8-12 Hz. For each frequency band, associations between fMRI and electrophysiological coherence were assessed using Spearman rank correlations across the eight condition-level observations defined by region pair × group × period.

### CAPs and C-mode analysis

Recurring fMRI states were identified using the co-activation mode (C-Mode) framework (Gutierrez-Barragan et al., 2019; Huang et al., 2020; Liu et al., 2013). This framework reduces spatially opposing coactivation patterns (CAPs) into coherent C-Modes. CAPs were mapped as previously described (Gutierrez-Barragan et al., 2024). Briefly, after processing, motion-contaminated rsfMRI volumes (framewise displacement > 0.075 mm) were removed, and the remaining rsfMRI volumes from all animals were pooled across groups and concatenated. CAPs were identified using k-means clustering, using spatial correlation as distance metric, 500 iterations and 5 random initializations. Clustering solutions ranging from k = 2 to 20 were evaluated across five independent runs. Group-level CAP maps were obtained by averaging all volumes assigned to a given cluster and converting voxel values to T-scores. Subject-level CAP maps were generated by averaging, within each subject, the frames assigned to each group-level cluster.

The optimal number of clusters was determined by evaluating explained variance and topographic stability across clustering solutions as previously described (Gutierrez-Barragan et al., 2024). Explained variance increased with k, reaching a plateau for k > 6 (Supplementary Figure 7a). CAP stability across increasing partitions was assessed using the Hungarian algorithm. Stable solutions were observed for k = 6–10. Within this range, we selected k = 10 because it provided the clearest segregation of opposing network states compared with lower-dimensional solutions used in previous mouse rsfMRI studies (Gutierrez-Barragan et al., 2019; Gutierrez-Barragan et al., 2024; Gutierrez-Barragan et al., 2022). Specifically, k = 10 yielded stronger CAP/anti-CAP spatial anticorrelation than k = 6 or k = 8 solutions (mean absolute pairwise spatial correlation: k = 10, |r| = 0.96; k = 8, |r| = 0.80; k = 6, |r| = 0.71; Supplementary Figure 7). This stronger spatial polarity supports the interpretation of the identified CAP/anti-CAP pairs as reciprocal expressions of coherent network modes, rather than independent transient activation patterns (Gutierrez-Barragan et al., 2024).

CAP pairs were next reduced to C-modes as previously described (Gutierrez-Barragan et al., 2024). For each CAP/anti-CAP pair, t-score-normalized maps were subtracted from one another, and the resulting difference map was divided by two to obtain a single mean C-mode map. To assess the network constituents of each C-mode, mean C-mode values were extracted from voxels located within predefined functional network masks. For each mouse, C-mode occurrence rate was computed as the proportion of frames assigned to either member of the corresponding CAP/anti-CAP pair. Because C-mode occurrence rates constitute compositional data, group differences in C-mode occurrence rates were assessed using multivariate analysis of variance (MANOVA) on isometric-log ratio (ILR)-transformed occurrence rates (Boogaart & Tolosana-Delgado, 2008).

To investigate transitions between dynamic states, CAP occurrence sequences were converted to C-mode labels. Transition counts were then pooled across subjects within each experimental group, without including transitions spanning subject boundaries, and used to estimate first-order Markov transition probability matrices. Only transitions within the same subject were included. Persistence probabilities were quantified from matrices including self-transitions, whereas transition probabilities between different C-modes were estimated after removing consecutive repetitions of the same C-mode to reduce autocorrelation driven by C-mode dwell times (Cornblath et al., 2020; Huang et al., 2020).

The entropy of Markov trajectories (HMT) was computed from the transition probability matrices as previously described (Ekroot & Cover, 1993; Gutierrez-Barragan et al., 2024; Huang et al., 2020). HMT quantifies the descriptive complexity of transition trajectories terminating in each target state, with higher values indicating more uncertain paths and, operationally, lower accessibility of that state. Group differences in persistence probabilities, transition probabilities, and HMT values were assessed using permutation testing. Null distributions were generated by randomly permuting group labels across subjects and recomputing the corresponding group-level metrics at each iteration. Multiple comparisons were controlled using the Benjamini–Hochberg false discovery rate procedure.

## Notes

### Competing Interest Statement

The authors have declared no competing interest.

