## Supplementary Figures for "Endogenous oxytocin reconfigures brainwide network dynamics toward salience-related states"

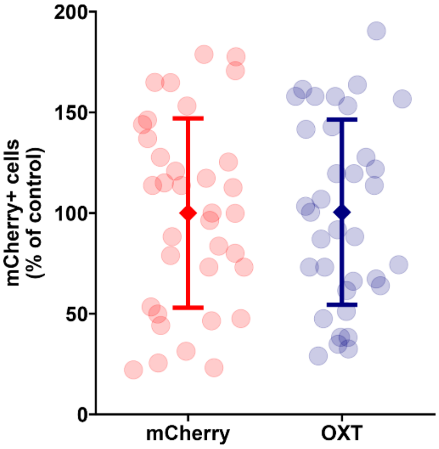


*Supplementary Figure 1. Comparison of hM3Dq-mCherry and Oxt-positive cell counts in the PVN. Quantification of hM3Dq-mCherry and OXT positive cells in the paraventricular nucleus on adjacent sections processed with immunohistochemistry and in situ hybridization, respectively (n=34 sections from 5 animals). Points represent positive cell counts per section, expressed as percentage of the mean Oxt-positive cell count. Error bars indicate standard deviation. Equivalence between RFP-positive and Oxt-positive cell counts was assessed using a mixed-effects two one-sided test (TOST) procedure, with predefined equivalence bounds of ±10% (mixed-effects TOST for equivalence: lower bound = −10%, p = 0.03; upper bound = 10%, p = 0.04).*

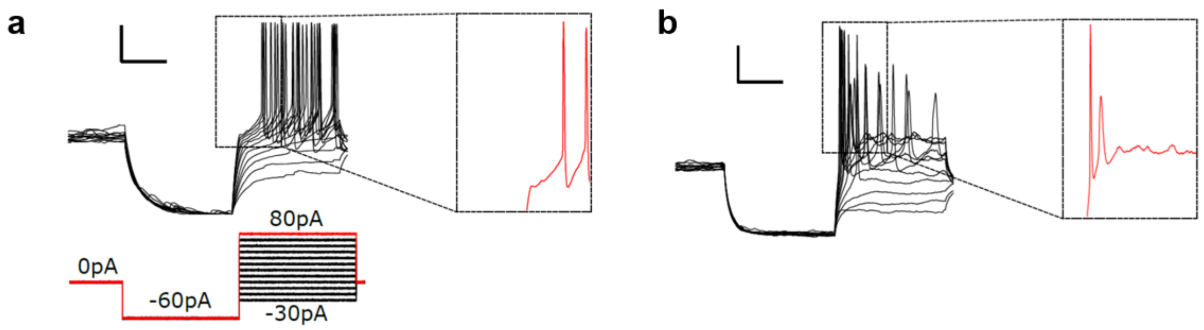


*Supplementary Figure 2. Electrophysiological characterization of PVN neurons in OXT-hM3Dq mice. (a) Traces showing electrophysiological identification of a magnocellular neuron in the paraventricular nucleus (PVN) during patch-clamp recordings under current-clamp mode displaying inward rectification, and strong transient outward rectification (black arrow in zoomed trace) in response to a series of depolarizing current pulses delivered at a hyperpolarized membrane potential. Scale bars are 10 mV and 100 ms.* ***(b)*** *Traces showing electrophysiological identification of a parvocellular neuron in the PVN during patch-clamp recordings under current clamp mode (I = 0 pA) displaying lack of transient outward rectification (black arrow in zoomed trace) in response to depolarizing current pulses delivered at a hyperpolarized membrane potential. Scale bars are 10 mV and 100 ms.*

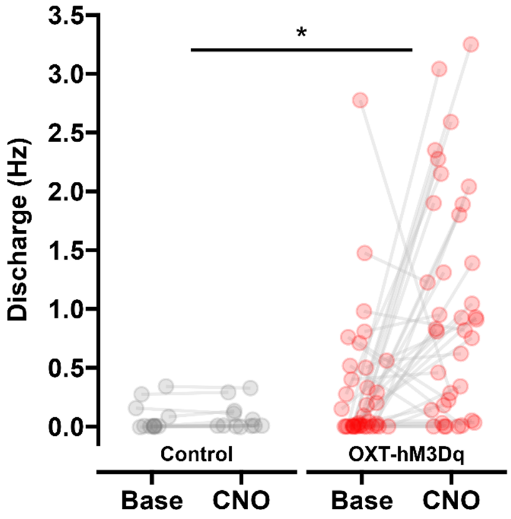


*Supplementary Figure 3. Electrophysiological characterization of all patched PVN neurons across groups. Whole-cell patch-clamp recordings were used to quantify firing frequency before and after CNO bath application in PVN neurons from OXT-hM3Dq and control mice. All recorded neurons were included in this analysis, irrespective of their electrophysiological profile, or responsiveness to CNO. As OXT-expressing neurons constitute only a subset of PVN neurons, this analysis may include cells lacking hM3Dq expression, and that as such were not expected to respond to CNO. CNO significantly increased firing frequency in OXT-hM3Dq mice compared to control littermates (treatment × genotype interaction, linear mixed model, permutation *$p < 0.05$). Points represent individual cells.*

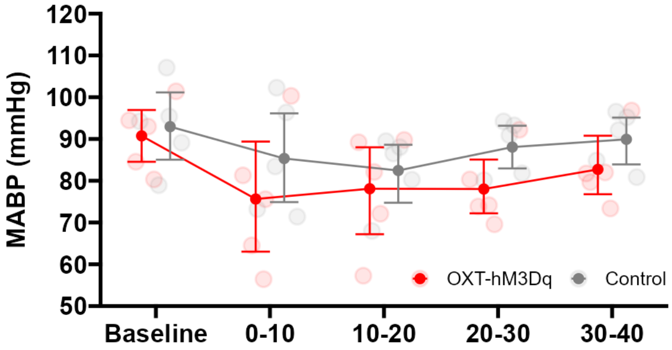


*Supplementary Figure 4. Arterial blood pressure after J60 administration.* *Mean arterial blood pressure (MABP) was recorded in male control (n = 5) and OXT-hM3Dq (n = 5) mice before and after administration of JHU37160 (1 mg/kg, i.p.; time = 0). Data show comparable mean arterial blood pressure between groups before and after J60 injection (mixed-effects TOST for equivalence compared to baseline, ΔL = -10, ΔU = 10; t=10-20: p<0.02; t=20-30: p<0.003; t=30-40: p<0.0005). Data are shown as individual subject values and group mean ± 95% CI.*

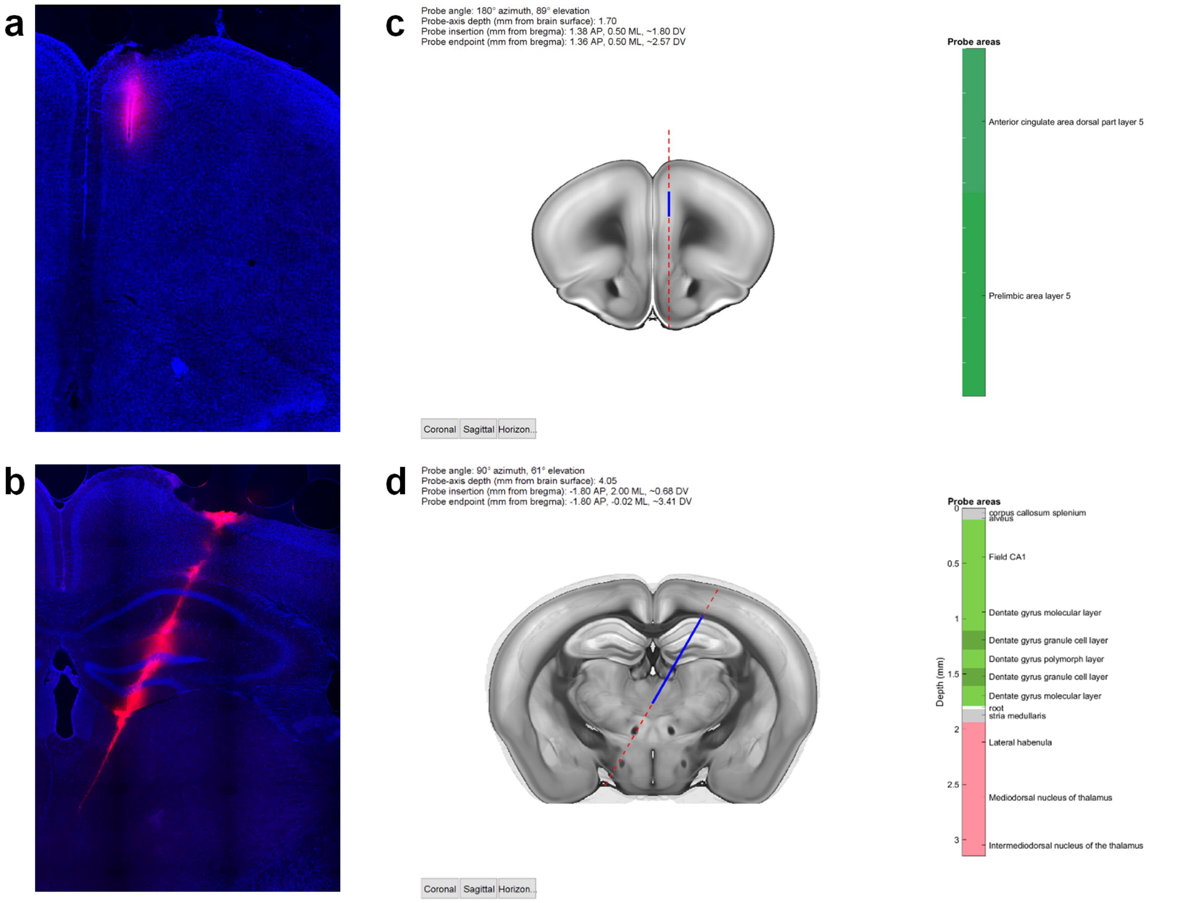


*Supplementary Figure 5. Electrode location.* *(**a-b**) Representative coronal sections showing DiI labeling (red) along probe tracks targeting the prefrontal cortex (PFC,* ***a****) and dorsal hippocampus and mediodorsal thalamus (dHPC/mdThal,* ***b****) Sections were counterstained with DAPI for anatomical localization. (**c-d**) Probe trajectories were reconstructed using the Neuropixels Trajectory Explorer. Using probe length, insertion point, insertion angle, and insertion depth, the anatomical location of each recording site was assigned according to the Allen Common Coordinate Framework. Region assignments were exported separately for each mouse to account for minor variability in probe placement and were used for subsequent analyses.*

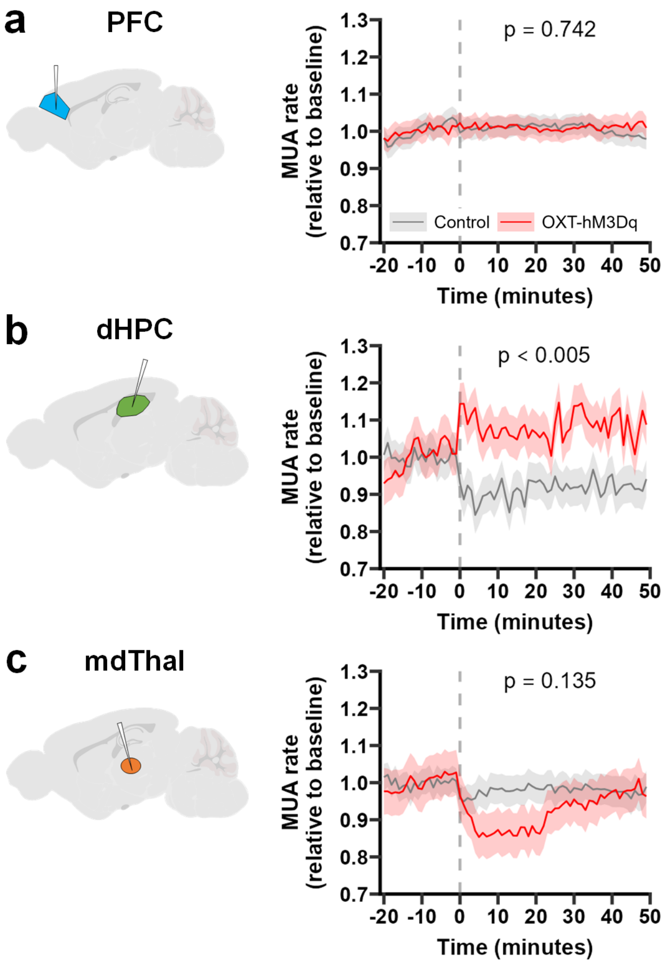


*Supplementary Figure 6. Local changes in multi-unit activity produced by endogenous OXT release. (a-c) Multi-unit activity (MUA) traces obtained in the medial prefrontal cortex (PFC, a), dorsal hippocampus (dHPC, b) and medial dorsal thalamus (mdThal) in OXT-hM3Dq (n = 9) and control (n = 11) male mice before and after administration of J60 (1 mg/kg, i.p.; time = 0). Solid lines show group means and shaded bands indicate bootstrapped 95% confidence intervals. Average values were computed for the pre- and post-injection periods, and group differences across periods were analyzed using a linear mixed-effects model with fixed effects of group, period, and their interaction, and random intercepts for channels nested within animals: firing rate ~ group × period + (1|animal/channel). P values indicate the significance of the group × period interaction.*

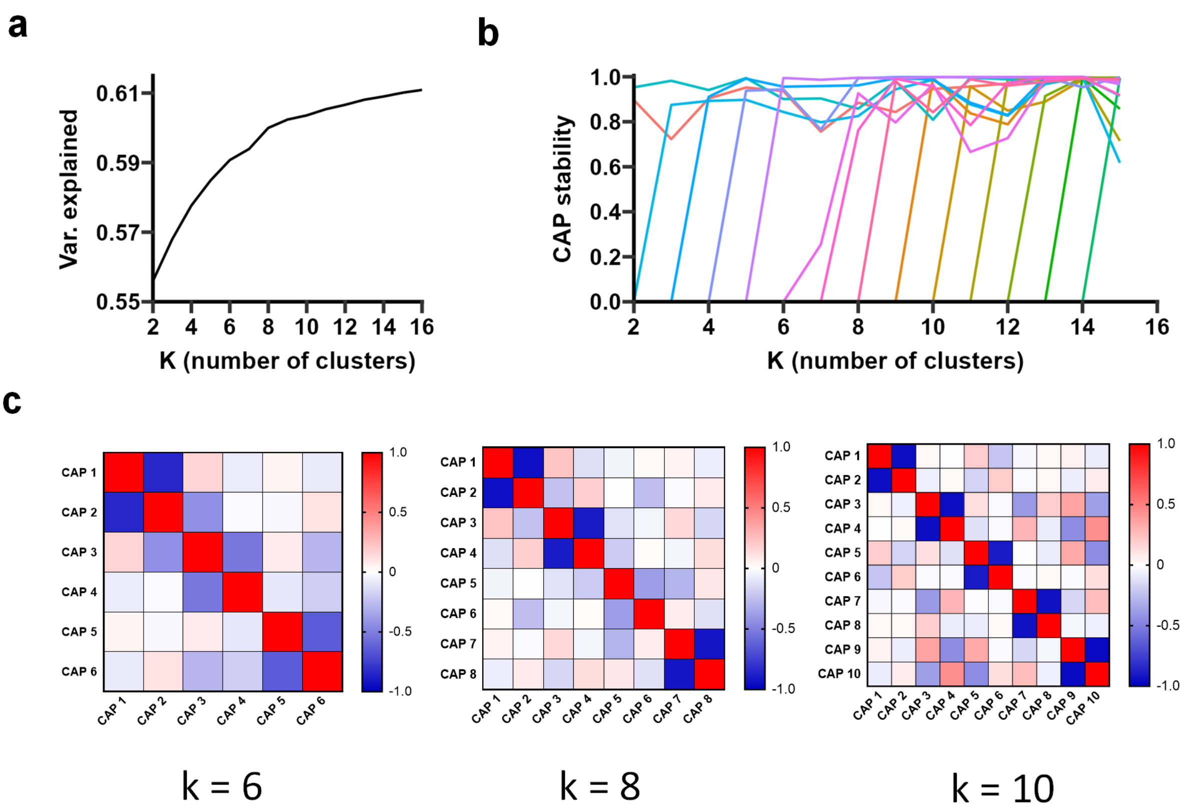


*Supplementary Figure 7. K-means clustering and C-mode number selection. (**a**) Variance explained by increasing numbers of clusters (2 < k < 16). A variance-based criterion identified k = 8 as the point beyond which additional clusters explained <1% additional variance. (**b**) CAP stability across increasing numbers of clusters. Stability was assessed by calculating the spatial correlation between CAPs from consecutive clustering solutions after optimal matching using the Hungarian algorithm. CAP stability for most CAPs dropped below 0.7 for k > 12. (**c**) Between-CAP similarity matrices showing spatial correlation between group-average CAP maps for k = 6, k = 8, and k = 10. The mean pairwise spatial correlation across matched CAP/anti-CAP pairs increased from |r| = 0.70 at k = 6 and |r| = 0.80 at k = 8 to |r| = 0.94 at k = 10. This stronger CAP/anti-CAP correspondence supported the selection of k = 10, yielding five pairs of opposing CAPs that defined the five C-modes analyzed in this study.*
